# Managing soil sustainably on small-scale vegetable farms: Lessons learned from high tunnel and open field vegetable production

**DOI:** 10.64898/2026.01.23.701315

**Authors:** Natalie Hoidal, Shane Bugeja, Julie Grossman, Adria Fernandez, Anna M. Cates, Kathryn M. LaBine, Devanshi Khokhani, Paulo Pagliari

## Abstract

Small-scale vegetable farms are increasingly important to local food systems, but the soils on these farms are not well understood, particularly in high tunnel production environments. Therefore, this study aimed to 1. Compare soil nutrients and soil health metrics in high tunnels and nearby open fields. 2. Document soil nutrient accumulation on diversified vegetable farms and assess loss potential. 3. Explore the impacts of specific management practices (input use, cover crops, tillage, and soil testing) and farm demographics on a variety of soil health and soil nutrient metrics. Just under half of the high tunnels in this study had soluble salt accumulation, which was associated with higher soil nitrate concentrations. The pH of many high tunnel soils was above the optimal range for crop production, which was correlated with irrigation water alkalinity. Some high tunnel soils had rapid water infiltration rates, with implications for irrigation water management. Both high tunnel and open field soil were rich in nutrients compared with other Minnesota farms. Preliminary assessments suggested risks to surface and groundwater from nutrient runoff and leaching. While farmer experience and more years in vegetable production were negatively associated with soil health metrics, management practices including reduced tillage, organic management, and application of plant-based compost were positively associated with soil health. Cation exchange capacity and permanganate oxidizable carbon did not provide significantly more insight than simply measuring organic matter. Arbuscular mycorrhizal fungal spore counts were inconclusive, but aggregate stability and bulk density were responsive to farmer reported soil management activities.

**Core ideas:**

- High tunnel soil tends to be rich in nutrients and organic matter. They also accumulate soluble salts, likely from excess inputs
- Irrigation water routinely tested high in pH and alkalinity. These factors may explain high soil pH in high tunnels.
- Small-scale vegetable farms often have high concentrations of soil nutrients in both high tunnels and open fields, with potential to cause environmental contamination through leaching and runoff
- Vegetable production may be inherently hard on soil health, but conservation practices including reduced tillage, organic management, and use of plant-based composts can improve soil health in these production systems

## 1 INTRODUCTION

Small-scale vegetable farming is becoming an increasingly important part of the food system in the Upper Midwest and across the United States. In Minnesota, the number of farms growing fresh market vegetables on a small scale has increased with each census cycle, with the largest increases among those growing on less than 10 acres, young farmers, and beginning farmers (United States, 2022). From 2012 to 2022, Minnesota gained 247 small-scale fresh market vegetable farmers (<50 acres) despite the state losing 9011 total farms during the same period (United States, 2012, 2022). This trend is consistent across the country (United States, 2022). Beginning farmers across the United States with limited access to land and capital are frequently drawn to vegetable production due to perceptions of fewer barriers to entry, and profitability at a small scale (DeLong et al., 2023). In Minnesota, farmers growing small-scale vegetables tend to be younger and more diverse than the average farmer growing field crops, with less formal training and mentorship compared to more established farmers (Bailey & Kagan, 2020; Hoidal et al., 2024; Sullivan, 2015). Developing a better understanding of soil health and nutrient management trends in these systems is important to ensure that farmers begin to use practices that will allow them to be successful and maintain sustainability.

Many fresh market vegetable farms utilize high tunnels. High tunnels are enclosed growing structures typically covered in 0.101 to 0.152 mm thick plastic, with limited to no supplemental heating and in-ground crop production. These structures extend the growing season in cool climates and generally provide more control of the growing environment (Lamont, 2009). In 2009 the Natural Resource Conservation Service (NRCS) launched the Seasonal High Tunnel Initiative to provide cost share funding and technical assistance to farmers to construct high tunnels as part of the Environmental Quality Incentives Program. It began as a three-year pilot program, and thereafter became an established conservation practice standard (standard #235; (National Sustainable Agriculture Coalition, 2016). To date, NRCS has funded 26,216 high tunnels nationally, including 694 in Minnesota (Pierre et al., 2024). As high tunnels have become more common, Extension educators in Minnesota report that farmers begin to observe significant challenges with productivity after a few years of high tunnel production. Farmers have expressed concerns about yield declines presumed to be due to salinity issues, disease pressure, and fertility challenges in high tunnels (Bruce et al., 2019; Pierre et al., 2024).

Heavy application of composted animal manure is relatively common in high tunnels, posing potential problems for nutrient and soluble salt accumulation (Pierre et al., 2024; Rudisill et al., 2015). In a study of 27 Pennsylvania high tunnels, 96% had phosphorus levels in excess of crop needs, sometimes as high as 10x the “exceeding crop needs” threshold (Sánchez & Ford, 2023a). The same study found that 48% of high tunnels had a pH above 7.0 (Sánchez & Ford, 2023b). However, studies on high tunnel soil dynamics have thus far been limited, and more research is needed to understand soil, nutrient, and water dynamics in these structures (Pierre et al., 2024), especially as compared to non-high tunnel vegetable production areas. Collectively, these factors may result in increased pressure on soil in high tunnels to sustain intensive vegetable production.

Some of these challenges may extend beyond high tunnels into open field production. Excess phosphorus accumulation has been documented in small-scale urban agriculture in Minnesota (Nicklay et al., 2019; Shrestha et al., 2020; Small et al., 2019) and noted anecdotally more broadly on rural and peri-urban vegetable farms by University of Minnesota Extension educators while providing technical assistance. A 2019 needs assessment (Klodd & Hoidal, 2019) found that 75% of small-scale fruit and vegetable farms in Minnesota are either certified organic or use exclusively organic practices. As such, many farms rely on compost, composted manure, or fresh manure as a source of nutrients for their crops (Hoidal, Bugeja, et al., 2025). There is significant confusion among beginning farmers about how to use compost effectively (Hoidal et al., 2024) and farmers tend to apply as much compost as possible to their farms, or apply set amounts (e.g. deep compost mulch systems), rather than applying compost and composted manure based on soil tests (Hoidal, Bugeja, et al., 2025). Reliance on these inputs to meet crop nitrogen needs tends to supply excess phosphorus and potassium (Möller, 2018) and regular use of compost can result in nitrogen applications in excess of crop needs (Ruch et al., 2023). Additionally, compost and manure have highly variable nutrient contents, and unless farmers are consistently testing inputs, averaging across reported values when calculating application rates can result in significant over or under application of nutrients (Wilson, 2021). Since most guidelines for nutrient management (e.g. Rosen & Eliason, 2005) are reported in pounds per acre, balancing crop nutrient needs at a smaller scale, especially when using organic inputs, is challenging at best. This challenge is exacerbated when farmers have a highly diversified portfolio of crops, each with different nutrient needs.

Based on these challenges, a study of 100 small-scale (<50 acres) diversified vegetable farms across Minnesota was conducted to develop a better baseline understanding of soils in both fields and high tunnels in these systems. The study had three primary objectives: 1. Compare nutrients and soil health metrics in high tunnels and nearby open fields. 2. Document soil nutrient accumulation on diversified vegetable farms and attempt to assess leaching or runoff risk. 3.

Explore the impacts of specific management practices (including input use, cover crops, tillage, and soil testing) and farmer demographics on a variety of soil health and soil nutrient metrics. The study team hypothesized that we would see high levels of phosphorus accumulation in both high tunnels and fields, salt and pH issues in high tunnels, and high tillage intensity in both environments.

## 2 Materials and Methods

### 2.1 Recruitment and survey

In the spring of 2023, a team of 16 Extension educators with the University of Minnesota visited 100 small-scale (<50 acres) vegetable farms to conduct soil tests in open fields and high tunnels, along with surveys. Farm selection criteria, recruitment methods, and survey methods are thoroughly documented in a previous manuscript (Hoidal, Bugeja, et al., 2025). The tests conducted and survey data collected at each farm are listed in Table 1 and described below.

**Table 1.** Tests conducted at each of the 100 farms.

| Soil physical properties | Soil chemical properties | Soil biological properties | Water tests | Survey data |
| --- | --- | --- | --- | --- |
| • Soil texture<br>• Bulk density<br>• Aggregate stability<br>• Infiltration rate | • pH<br>• Phosphorus (Bray-1 and Olsen) + total phosphorus<br>• Potassium<br>• Nitrate nitrogen<br>• Electrical conductivity<br>• Cation exchange capacity | • Organic matter<br>• Permanganate oxidizable carbon (POX-C)<br>• Arbuscular mycorrhizal fungi spores | • pH<br>• Alkalinity | • Years of experience (farmer)<br>• Years in production (site)<br>• Farm size<br>• Organic status<br>• Frequency of cover crop use<br>• Tillage intensity<br>• Type and frequency of inputs |

### 2.2 In-situ soil analysis

Water infiltration rate was measured twice *in situ* at each farm using a single ring (Natural Resources Conservation Service, 2001). A 7.62 cm diameter metal cylinder was hammered 7.62 cm into the soil using a rubber mallet and covered in plastic wrap. A volume of 107 ml of water was poured into the plastic-lined ring. Infiltration was timed from pulling the plastic wrap away to the disappearance of ponded moisture at the surface. This amount of water mimics 25.4 mm, or one inch of rainfall for the volume of the cylinder. This was repeated twice in each cylinder, totaling 50.7 mm of infiltration water. The procedure was stopped if the time needed for the water to infiltrate exceeded 10 min for the first 25.4 mm of water, or after 5 min for the second 25.4 mm of water. Data was originally collected as # seconds to infiltrate 25.4 mm water, then converted to mm infiltration per hour by converting seconds to minutes, then calculating inches per hour using Equation 1.

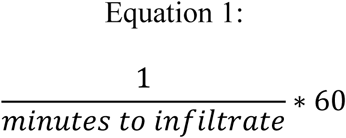

Finally, inches per hour was converted to mm per hour. Spring of 2023 was generally wet, and so field conditions were often saturated, but soil conditions varied considerably from site to site since samples were collected over a six-week period.

### 2.3 Laboratory soil analysis

Soil samples were collected with a 2.54-cm metal soil probe from a 0-15 cm depth interval after pushing any surface mulch to the side. Between 15 to 20 subsamples were collected from each site (field or high tunnel) into a single five-gallon container, mixed by hand, and aggregated into a single composite sample for each field or high tunnel. Approximately 1 liter of soil was collected from each site and allowed to air dry. The composite samples from each location were sent to the University of Minnesota Research Analytical Laboratory to analyze pH, Bray-1 phosphorus, Olsen phosphorus (for all soils with a pH greater than 7.4), exchangeable potassium, nitrate, electrical conductivity (1:1 suspension method), cation exchange capacity (CEC; summation method), and organic matter (OM) (University of Minnesota, 2024b, 2024a). Since the 1:1 suspension method is typically just used for screening electrical conductivity, all samples with an electrical conductivity measurement above 0.5 mmhos cm^-1^ were re-screened using a saturated paste extract (University of Minnesota, 2024b). All statistical analyses for electrical conductivity were performed on the subset of data using the saturated paste extract.

Soil particle size distribution was determined by laser particle size analysis, with quality control samples determined by the pipette and hydrometer methods (Grigal, 1973; Soil Survey Staff, 2022). Samples for particle size analysis were air-dried, passed through a 2 mm sieve, and hand homogenized prior to subsampling. Three 0.5 g subsamples were then obtained and dispersed overnight in 200 ml of de-ionized water with Na-hexametaphosphate (5 ml) and Na-hypochlorite (5 ml) prior to standard analysis on a Malvern Mastersizer 3000. The refractive indices of soil and the water-based dispersant were assumed to be 1.549 and 1.33, respectively (Miller & Schaetzl, 2012). Extensive in-house validation was conducted on known size fractions of sand, silt and clay to optimize laser particle size data for comparison with the more traditional hydrometer method. Based on this in-house compilation, the clay-silt break was set at 8 µm, which is consistent with other studies (Konert & Vandenberghe, 1997). To eliminate bias due to small subsample sizes, triplicate analyses of each of the three subsamples were conducted and compared. The set of three that was most different from the other (two) subsamples was subsequently discarded. The two most comparable triplicate sets were then used to calculate the mean particle size distribution (n = 6) for use in subsequent analyses. For each set of 20 samples, a standard (ISO 12103-1, A2 FINE TEST DUST, Powder Technology, Inc.) was run to ensure internal quality control standards were met.

At each site, soil was collected using a 7.62 cm diameter metal ring, pounding it 7.62 cm into the soil, and carefully collecting the entire contents of the core. The soil from each core was oven dried at 32.2°C and weighed to determine the bulk density in g/cm^3^.

Aggregate stability samples were taken using a trowel to carefully extract a cylinder of soil approximately 15 cm deep. These samples were broken by hand along natural planes of weakness and dried at 35°C in a paper bag. Aggregate stability was measured by a modified wet sieving process (Yoder, 1936). Four sieves in a PVC cylinder were used to separate the sample into four aggregate size classes: >2 mm, 1-2 mm, 0.25 mm-1 mm, 0.053-0.25 mm. Fifty grams of sample were spread evenly on top of the 2 mm sieve, and the cylinder was filled with deionized water up to the top sieve surface, allowing 10 minutes of capillary wetting. Then stacked sieves were mechanically raised and lowered at a rate of 25 rpm for 10 mins. Soil remaining on top of the four sieves was rinsed into a beaker and oven dried at 50-60°C until completely dried. Any sample that fell through the 0.053 mm sieve was estimated by subtraction from the moisture-corrected starting mass. Sand corrections were done to the >2 mm and 1-2 mm samples.

Permanganate oxidizable carbon (POX-C) was assessed according to Weil et al. (2003) Briefly, 1.25 g of air-dry soil was reacted with a 0.2 M KMnO_4_ solution, which is a strong oxidizer. Diluted supernatant from each sample was transferred to 96-well plates and measured for absorbance at 540 nm on a spectrophotometer. Absorbance was fitted to a standard curve using standards with known concentration made with KMnO4 and calculated to determine C oxidation by KMnO_4_ reaction.

Arbuscular mycorrhizal fungal spores were extracted and enumerated using the wet sieving method (Gerdemann & Nicolson, 1963), decanting, and followed by sucrose density gradient centrifugation. Briefly, 50 g of a soil sample was placed in a Waring Blender and blended at high speed for approximately 5 seconds to break up root fragments and release spores attached to roots or soil aggregates. The blended material was immediately poured through two sieves - the top sieve (opening of 500 µm), which captures roots and large debris, and the bottom sieve (opening of 38 µm), which captures the majority of spores. The material on the bottom sieve was collected in a 50-mL beaker with a rubber policeman and then transferred into 50-mL tubes containing a 60% sucrose solution and water. These tubes were centrifuged (960 x g) for 5 minutes in a swinging-bucket rotor in a tabletop centrifuge. At the end of the run, the sucrose layer where the spores were suspended was collected by pouring it onto a 38 µm sieve. These spores were washed with water before being collected on the petri plates with the help of a rubber policeman. The bottoms of the plates were labeled with blue horizontal and red vertical lines to count the spores per square under the stereo microscope and the number of spores were reported per 50 g of soil sample.

One liter of water was collected from each site from the irrigation source after running the water for 30 seconds to clear debris. The water samples were also sent to the RAL for pH and alkalinity tests (University of Minnesota, 2024b).

### 2.4 GIS analysis & nutrient loss risk screening

Using ArcGIS, the distance and slope from each site (field or high tunnel) to the nearest surface water body (including lakes, wetlands, and streams) was calculated following hydrologic flow paths, and reported in meters. This was done using the National Wetland Inventory for Minnesota (Minnesota Department of Natural Resources, 2019), and the Lidar Elevation Data for Minnesota: 2008-2012 (Minnesota Geospatial Information Office, 2012). The depth to the water table from each site was also determined using the shapefile: Water-Table Elevation and Depth to Water Table, Minnesota Hydrogeology Atlas series HG-03 (Minnesota Department of Natural Resources, 2016).

Distance data was then compared with distance-based phosphorus runoff risk metrics (Sharpley et al., 2003). A rough phosphorus loss screening was applied to each sample using the Pennsylvania Phosphorus Index screening tool (Sharpley et al., 2003). We selected this over the Minnesota Phosphorus Index screening due to its simplicity and because the Pennsylvania index aligned closely with our outcomes of interest. The Pennsylvania screening tool uses the Mehlich-3 method for determining soil phosphorus, and so we used Equation 2 to convert Bray-P1 concentrations to Mehlich-3 (Culman et al., 2019).

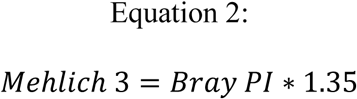

To assess risk of nitrogen loss, soil texture was reclassified from 11 texture groups into the four simplified hydrologic soil groups (National Oceanic and Atmospheric Administration, 2022). This was then combined with depth to water table. Soils in groups A and B with a water table depth of 0-10 m were characterized as high and moderate infiltration risk, respectively. Based on the Minnesota Department of Agriculture’s Vulnerable Groundwater Area Map, karst geology and shallow bedrock were added as risk factors (Minnesota Department of Agriculture, 2025).

Using ArcGIS, farms were mapped over shapefiles indicating karst geology and shallow bedrock, and farms with these features were flagged, and nitrogen loss risk was elevated (Figure 1).

**Figure 1.**
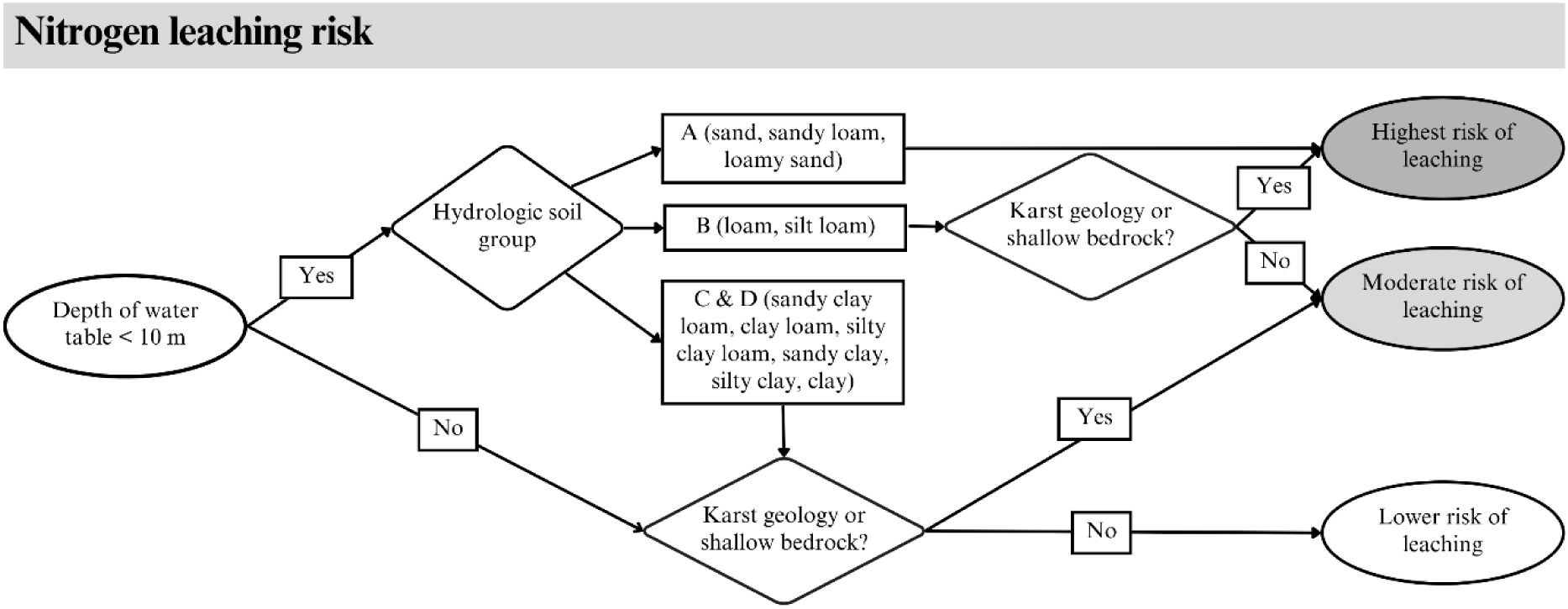
Flow chart indicating the process for determining nitrogen leaching risk.

### 2.5 Comparisons to statewide data

Data from small-scale vegetable farms in this study were compared to statewide farm averages utilizing an unpublished dataset from the University of Minnesota’s Research Analytical Laboratory. This data includes 22,826 farm and field soil tests conducted at the lab between 1998 - 2016. Not every soil tests included each variable measured in this paper; as such the n value is cited for each variable.

### 2.6 Statistical analysis

All data was analyzed using R (v4.2.2; R Core Team, 2022). Graphs were generated with the ggplot2 package, and summary statistics were generated using basic commands within the dplyr package. Welch’s two-sample t-test was used to assess differences between high tunnels and fields for individual variables. Spearman’s rank correlation coefficient was used to measure relationships between ranked categorical variables and dependent continuous variables. Ranks were assigned as follows: Farmer experience: farmers with <5 years experience < Farmers with 5-10 years experience < Farmers with 10+ years experience. Organic certification: Conventional < Using mostly organic practices < Not certified but using exclusively organic practices < Certified organic. Soil test frequency: I have never tested the soil < I tested it once when I started but have not done it again < Every 4+ years < Every 2-3 years < Every year. Cover crop frequency: never < Occasionally < Every 2-3 years <Every year. Input use: Never < Every few years < About once per year < More than once per year. Finally, Pearson’s correlation coefficient was used to measure relationships between continuous variables.

## 3 Results

### 3.1 High tunnel vs. field soils

Soil organic matter, CEC, and POX-C were all higher in high tunnels than in fields (Table 2), but in both environments, soil organic matter levels were considered “high” (Rosen & Eliason, 2005). All measured plant nutrients, including phosphorus, potassium, nitrate, ammonium, calcium, and magnesium, were higher in high tunnels than fields (Table 2).

**Table 2.** Soil test results from high tunnels and open fields at 100 vegetable farms in Minnesota. Means and standard deviations are reported for each variable, along with Welch’s two sample t-test results comparing field vs. high tunnel soils.

|  | High tunnel | Open field | Tunnel vs. field t-test |
| --- | --- | --- | --- |
| Soil organic matter (%) | $6.34 \pm 3.23$ | $4.66 \pm 2.35$ | $t(179) = -4.18, p < 0.001$ |
| Permanganate oxidizable carbon (mg C oxidized per kg dry soil) | $1248.97 \pm 490.60$ | $1006.41 \pm 531.77$ | $t(193) = -3.31, p = 0.001$ |
| Cation exchange capacity (cmol/kg) | $28.12 \pm 12.06$ | $20.44 \pm 12.42$ | $t(197) = -4.43, p < 0.001$ |
| pH | $7.13 \pm 0.56$ | $6.83 \pm 0.65$ | $t(193) = -3.46, p < 0.001$ |
| Electrical conductivity (1:1 suspension; mmhos $\text{cm}^{-1}$) | $0.69 \pm 0.48$ | $0.25 \pm 0.15$ | $t(116) = -8.78, p < 0.001$ |
| Phosphorus (Bray-P1; ppm) | $177.90 \pm 141.67$ | $88.94 \pm 70.87$ | $t(93) = -4.73, p < 0.001$ |
| Phosphorus (Olsen; ppm) | $101.66 \pm 73.07$ | $60.73 \pm 61.58$ | $t(31) = -2.03, p = 0.051$ |
| Potassium (ppm) | $328.28 \pm 318.85$ | $228.06 \pm 184.45$ | $t(157) = -2.71, p = 0.007$ |
| Nitrate (top 6” only; ppm) | $62.45 \pm 68.76$ | $11.69 \pm 10.22$ | $t(102) = -7.27, p < 0.001$ |
| Ammonium (ppm) | $8.53 \pm 9.21$ | $6.72 \pm 4.95$ | $t(150) = -1.72, p = 0.088$ |
| Calcium (ppm) | $4171.08 \pm 1787$ | $3170.32 \pm 1881.81$ | $t(197) = -3.86, p < 0.001$ |
| Magnesium (ppm) | $688.17 \pm 353.07$ | $450.77 \pm 345.28$ | $t(197) = -4.79, p < 0.001$ |
| Bulk density ($\text{g}/\text{cm}^3$) | $1.04 \pm 0.27$ | $1.14 \pm 0.29$ | $t(196) = 2.42, p = 0.165$ |
| **Proportion large soil aggregates** | 0.56 ± 0.25 | 0.55 ± 0.28 | t(186) = -0.35, p=0.728 |
| **AMF spore density (per 50 g of soil)** | 591.32 ± 389.42 | 526.74 ± 370.82 | t(192) = -1.10, p=0.236 |

Salt accumulation, as measured by electrical conductivity, was higher in high tunnels than in fields (Table 2), with a white crusting frequently observed on the soil surface. Electrical conductivity screening tests using the 1:1 suspension method showed soluble salt levels to generally be low, measured at <2 mmhos cm^-1^ in 100% of fields and 98% of tunnels (Rosen & Eliason, 2005). However, after re-screening all samples with a 1:1 suspension concentration of 0.5 mmhos cm^-1^ or higher using the saturated paste method, 26 of the tunnels were slightly saline (2-4 mmhos cm^-1^), and 15 were moderately saline (2-4 mmhos cm^-1^). The maximum electrical conductivity was 7.3 mmhos cm^-1^. Only one open field was very slightly saline. Higher soil nitrate was positively correlated to electrical conductivity (rho = 0.36; p-value 0.0196), whereas increased tillage was negatively correlated with electrical conductivity (rho = -0.38; p-value 0.0194). None of the following factors were correlated with electrical conductivity: farmer experience, organic certification, cover crop frequency, soil test frequency, age of field / tunnel, soil pH, water pH, water alkalinity, soil calcium, or application frequency of fresh manure, composted manure, vegetative compost, synthetic nitrogen, and synthetic fertilizers (Tables 4, 5, & 6).

**Table 3.** Infiltration rates for 25.4 mm and 50.7 mm of water applied to field and high tunnel soils. Means and standard deviations are reported for each variable, along with Welch’s two sample t-test comparing field vs. high tunnel soils.

|  | Field | High tunnel | Tunnel vs. field t-test |
| --- | --- | --- | --- |
| Infiltration rate (mm/hr) for first 25.4 mm / 1 inch of water | 2849 ± 8492 | 4349 ± 8475 | t(163) = -0.98, p=0.3286 |
| Infiltration rate (mm/hr) for first 25.4 mm / 1 inch of water | 3065 ± 9644 | 4465 ± 8673 | t(163) = -1.14, p= 0.258 |

**Table 4.** Distance from field and high tunnel sites to the nearest surface water (lake, river, wetland) calculated along hydrological flow paths, categorized by risk level of P losses to water outlined in and Sharpley et al. (2003).

|  | 0<br>(Lowest risk)<br>(>152 m) | 2<br>(152 -107 m) | 4<br>(76 - 107 m) | 6<br>(76 - 46 m) | 8<br>(Highest risk)<br>(<46 m) |
| --- | --- | --- | --- | --- | --- |
| # fields | 50 | 5 | 10 | 14 | 21 |
| # tunnels | 46 | 7 | 11 | 8 | 28 |

**Table 5.** Correlation coefficients and p-values from Spearman’s Rho test comparing ranked categorical variables with soil health metrics. Cells with shading represent significant correlations.

|  | Farmer experience | Organic certification | Soil test frequency | Frequency of cover crop use |
| --- | --- | --- | --- | --- |
| Soil organic matter (%) | -0.21<br>p = 0.0035 | 0.32<br>p = 0.0000 | -0.01<br>p = 0.9040 | -0.17<br>p = 0.0261 |
| Cation exchange capacity (cmol/kg) | -0.16<br>p = 0.0285 | 0.37<br>p = 0.0000 | -0.05<br>p = 0.5583 | -0.11<br>p = 0.1516 |
| Permanganate oxidizable carbon (mg C oxidized per kg dry soil) | -0.22<br>p = 0.0020 | 0.25<br>p = 0.0009 | -0.007<br>p = 0.9277 | -0.23<br>p = 0.0025 |
| Electrical conductivity (1:1 suspension; mmhos cm<sup>-1</sup>) | -0.03<br>p = 0.7255 | 0.17<br>p = 0.0287 | -0.01<br>p = 0.9226 | -0.14<br>p = 0.0613 |
| Phosphorus (Bray-P1; ppm) | 0.18<br>p = 0.0115 | -0.05<br>p = 0.5402 | 0.06<br>p = 0.4717 | -0.20<br>p = 0.0101 |
| Nitrate (top 6" only; ppm) | 0.05<br>p = 0.5295 | 0.10<br>p = 0.1758 | -0.06<br>p = 0.4486 | -0.15<br>p = 0.0512 |
| Bulk density (g/cm<sup>3</sup>) | 0.15<br>p = 0.0366 | -0.19<br>p = 0.0150 | -0.02<br>p = 0.8406 | 0.11<br>p = 0.1731 |
| % large soil aggregates | -0.23<br>p = 0.0015 | 0.12<br>p = 0.1309 | -0.13<br>p = 0.0981 | -0.12<br>p = 0.1409 |
| Infiltration rate (mm/hr) | -0.4<br>p = 0.6461 | 0.10<br>p = 0.2093 | 0.09<br>p = 0.2629 | -0.18<br>p = 0.0304 |
| AMF spore density (per 50 g of soil) | 0.11<br>p = 0.1348 | -0.26<br>p = 0.0007 | -0.07<br>p = 0.4009 | -0.10<br>p = 0.2181 |

The pH was also higher in high tunnels than in fields (Table 2), with the mean high tunnel pH just above the optimal range for vegetable production, 5.8-7.0 (Rosen & Eliason, 2005). It did not increase over time, with younger tunnels being just as likely to have a high soil pH as older tunnels (rho = 0.15, p-value = 0.1727) (Figure 1). More frequent use of synthetic nitrogen (rho = -0.27; p-value 0.0226) and synthetic fertilizers (rho = -0.28; p-value 0.0155) was associated with a lower soil pH. Higher soil calcium (rho = 0.48; p-value 0.0000) and higher irrigation water alkalinity (rho = 0.33; p-value 0.0007) were associated with a higher soil pH. In the subset of high electrical conductivity soils (those re-screened with the saturated paste extract), there was no correlation between electrical conductivity and pH (rho = 0.05; p-value = 0.7629).

Bulk density, the percentage of soil aggregates classified as “large”, and the number of arbuscular mycorrhizal fungal spores did not differ between high tunnel versus open field environments (Table 2).

### 3.2 Irrigation & soil water dynamics

The median irrigation water pH was 7.9 (s.d. 0.35), and the median alkalinity was 206 (s.d. 115.55). Both metrics are well above the ideal range of 5.0 - 7.0 for irrigation water pH, and <100 ppm for alkalinity (Cox, 1995). Only one farm had irrigation water with a pH less than 7.0, and only 10 farms had an alkalinity reading of less than 100 ppm.

High tunnel water infiltration rates were faster than fields (Table 3). Some tunnels drained very quickly, with 26% of high tunnel soils absorbing the first 25.4 mm of water in less than 30 seconds, and 21% percent absorbing the second 25.4 mm in less than 30 seconds. Educators who conducted infiltration tests with very fast absorption (i.e. less than 30 seconds) noted that the water passed so quickly through these soils that the soil remained dry at the surface after the infiltration test.

While the survey did not include questions about winter irrigation, educators noted that many of the high tunnels were empty and that the soil had been left bare and allowed to dry over the winter.

### 3.3 Nutrient accumulation

In both high tunnels and fields, median nutrient concentrations were above, and often well above the “very high” thresholds set for Minnesota vegetable soils (Rosen & Eliason, 2005). This was particularly notable for phosphorus, with a mean of 89 ppm (Bray-1) in open fields and 178 ppm (Bray-1) in high tunnels (Table 2). “Very high” thresholds for phosphorus are 41 ppm (Bray-1) (Rosen & Eliason, 2005). Of the 100 tunnels and 100 fields we sampled, 87% of tunnels and 84% of fields exceeded the “very high” threshold for soil phosphorus, and the average vegetable farm in our study had phosphorus levels well above the statewide farm median of 38.39 ppm (Bray-1) (Unpublished RAL data, n = 58983).

The average field soil had 11.69 ppm nitrate in the top 15 cm, equating to a 23.38 lbs / acre NO_3_-N credit in the top 15 cm, and the average high tunnel soil had 62.45 ppm nitrate, translating to a 124.9 lbs / acre NO_3_-N credit in the top 15 cm (Camberato & Nielsen, 2017). For a tomato crop, this amount of nitrate in the top 15 cm alone would account for 18-30% of recommended nitrogen fertility in fields, depending on soil type (Rosen & Eliason, 2005) and in a high tunnel, this would account for 36-59% of recommended nitrogen fertility depending on yield goal and soil type (Hoidal, Rosen, et al., 2025).

Calcium concentrations in open fields were more than twice as high as the statewide median calcium concentration for Minnesota farms (1414 ppm; RAL unpublished data, n=9999), and over three times higher in tunnels (Table 2). Additional calcium inputs are not recommended when soil calcium levels exceed 300 ppm (Rosen & Eliason, 2005).

### 3.4 Runoff and leaching risk assessment

Based on the screening tool used in the Pennsylvania phosphorus index (soil test P > 200 ppm according to Mehlich-3 and distance to water source <45.72 m), six fields and 15 high tunnels had a high enough phosphorus loss risk to be considered for a full phosphorus index assessment (Sharpley et al., 2003). None of the fields or high tunnels that met both criteria had soils in hydrologic group D (poorly drained soils with slow infiltration rates and high runoff risk). Using the same system, 50% of fields and 46% of high tunnels were located at a low-risk distance to water, whereas 19% of fields and 14% of tunnels were at a high-risk distance, and 21% of fields and 28% of tunnels were at a very high-risk distance to surface water (Table 4). A full phosphorus index analysis was not within the scope of this project.

Nearly two thirds of fields and high tunnels had a high to moderate risk of nitrate leaching based on soil type and water table depth: Of the 30 fields and 37 high tunnels with soils in hydrologic group A (sandy with high infiltration rates), 21 fields and 26 high tunnels had shallow (<10m) groundwater, representing the highest risk for leaching. Of the 53 fields and 58 high tunnels with soils in hydrologic group B (moderate infiltration rates), 39 fields and 38 high tunnels had shallow (<10m) groundwater, representing moderate risk for leaching. Among these high to moderate leaching risk soils, six fields and five high tunnels had an additional elevated risk due to karst geology, and one farm (both their field and high tunnel) had an elevated risk due to shallow bedrock. Among the soils classified as lower risk due to either clay soils or a deeper water table, six fields and four tunnels had karst geology, and three fields and three tunnels had shallow bedrock, elevating them to a potentially higher risk category. With the inclusion of karst geology and shallow bedrock as risk factors, over two thirds of the fields and high tunnels had a high to moderate risk of nitrate leaching.

### 3.5 Soil health practices and metrics

#### 3.5.1 Farmer demographics

Farmers with more years of experience had lower OM, CEC, POX-C, and aggregate stability (as measured by % very large aggregates) (Table 5). More experienced farmers also had more soil phosphorus and more compaction (Table 5). Similarly, fields that had been in vegetable production longer had more compact soil, slower infiltration, and lower aggregate stability (Table 6).

**Table 6.** Correlation coefficients and p-values from Pearson’s correlation coefficient. Tillage score calculation from (Hoidal et al., 2025). Cells with shading represent significant correlations.

|  | Tillage score | Years in vegetable production | Farm size (acres) |
| --- | --- | --- | --- |
| Soil organic matter (%) | -0.20<br>p = 0.0070 | -0.13<br>p = 0.1027 | -0.14<br>p = 0.0623 |
| Cation exchange capacity (cmol/kg) | -0.13<br>p = 0.0739 | -0.12<br>p = 0.1126 | -0.14<br>p = 0.0513 |
| Permanganate oxidizable carbon (mg C oxidized per kg dry soil) | -0.14<br>p = 0.0552 | -0.13<br>p = 0.0930 | -0.15<br>p = 0.0497 |
| **Electrical conductivity (1:1 suspension; mmhos cm<sup>-1</sup>)** | -0.19<br>p = 0.0105 | -0.09<br>p = 0.2651 | -0.06<br>p = 0.3860 |
| **Phosphorus (Bray-PI; ppm)** | -0.06<br>p = 0.4481 | 0.01<br>p = 0.9397 | -0.01<br>p = 0.8882 |
| **Nitrate (top 6" only; ppm)** | -0.16<br>p = 0.0278 | -0.06<br>p = 0.4734 | -0.04<br>p = 0.5672 |
| **Bulk density (g/cm<sup>3</sup>)** | 0.12<br>p = 0.0978 | 0.22<br>p = 0.0049 | 0.10<br>p = 0.1924 |
| **% large soil aggregates** | -0.20<br>p = 0.0058 | -0.21<br>p = 0.0075 | -0.14<br>p = 0.0712 |
| **Infiltration rate (mm/hr)** | -0.09<br>p = 0.2252 | -0.09<br>p = 0.2673 | -0.04<br>p = 0.6356 |
| **AMF spore density (per 50 g of soil)** | 0.09<br>p = 0.2458 | -0.07<br>p = 0.3927 | 0.14<br>p = 0.0640 |

Certified organic farmers and those using more organic practices had more OM, higher CEC and POX-C, less compaction, and higher soluble salts than conventional farms (Table 5). Farm size (measured in total farm acres in vegetable production) was negatively correlated with POX-C and had a weak negative correlation with OM and CEC. Farm size was not significantly correlated with other soil health metrics (Table 6).

#### 3.5.2 Farmer practices

Frequency of cover crop use was not correlated with tillage (X^2^ = 64.421, p = 0.9116), nor was it significantly correlated with farmer experience level (X^2^ = 5.287, p = 0.5057). More frequent cover crop use was associated with lower OM, slower water infiltration, and less phosphorus compared to less frequent users of cover crops (Table 5). When separated by production environment, the correlation between cover crop frequency and OM was more pronounced in open fields than in high tunnels (Figure 2). However, when the dataset was filtered to farmers with more than 10 years of experience, this trend became less clear (Figure 3).

**Figure 2.**
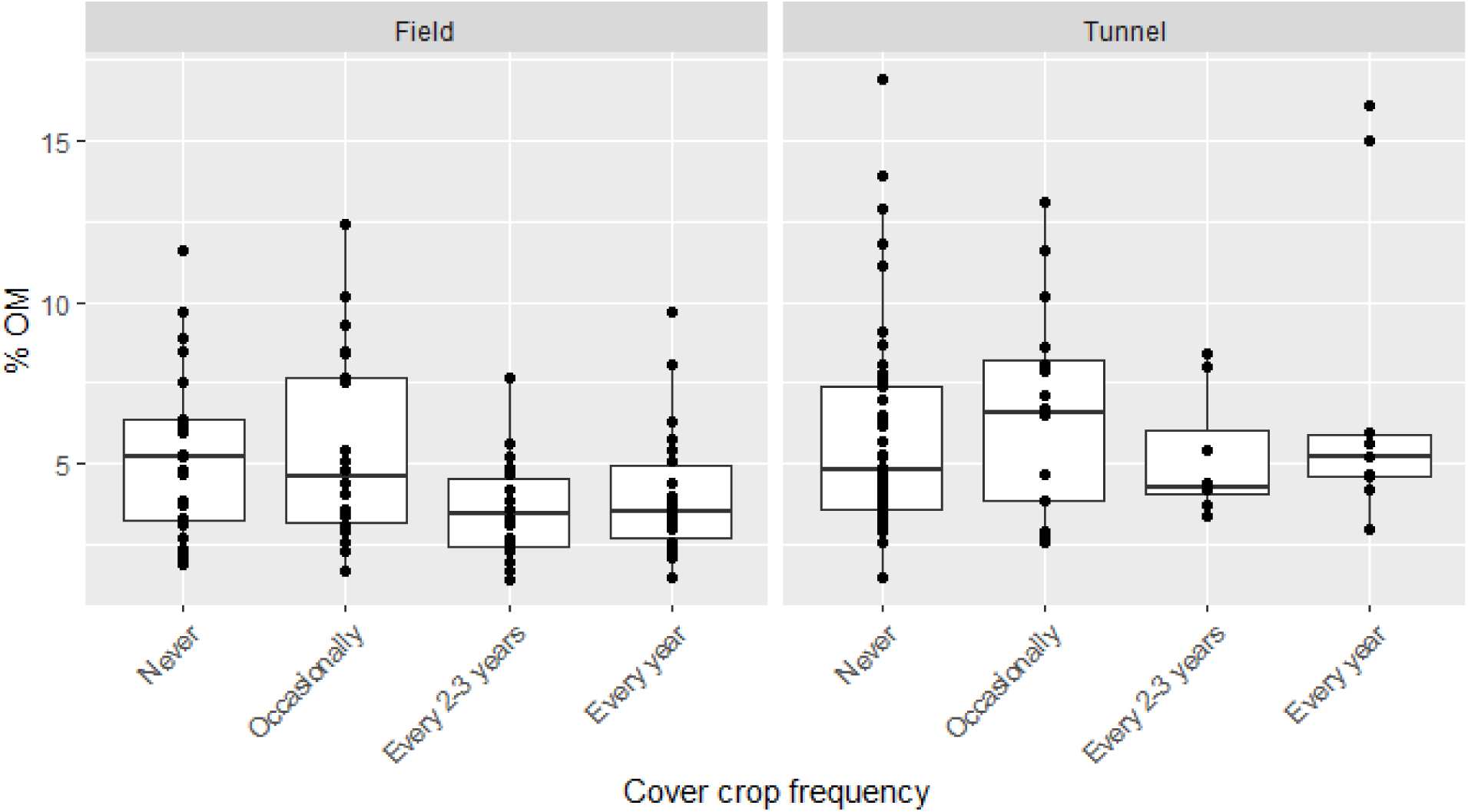
Cover crop frequency and its impact on soil organic matter in fields and high tunnels.

**Figure 3.**
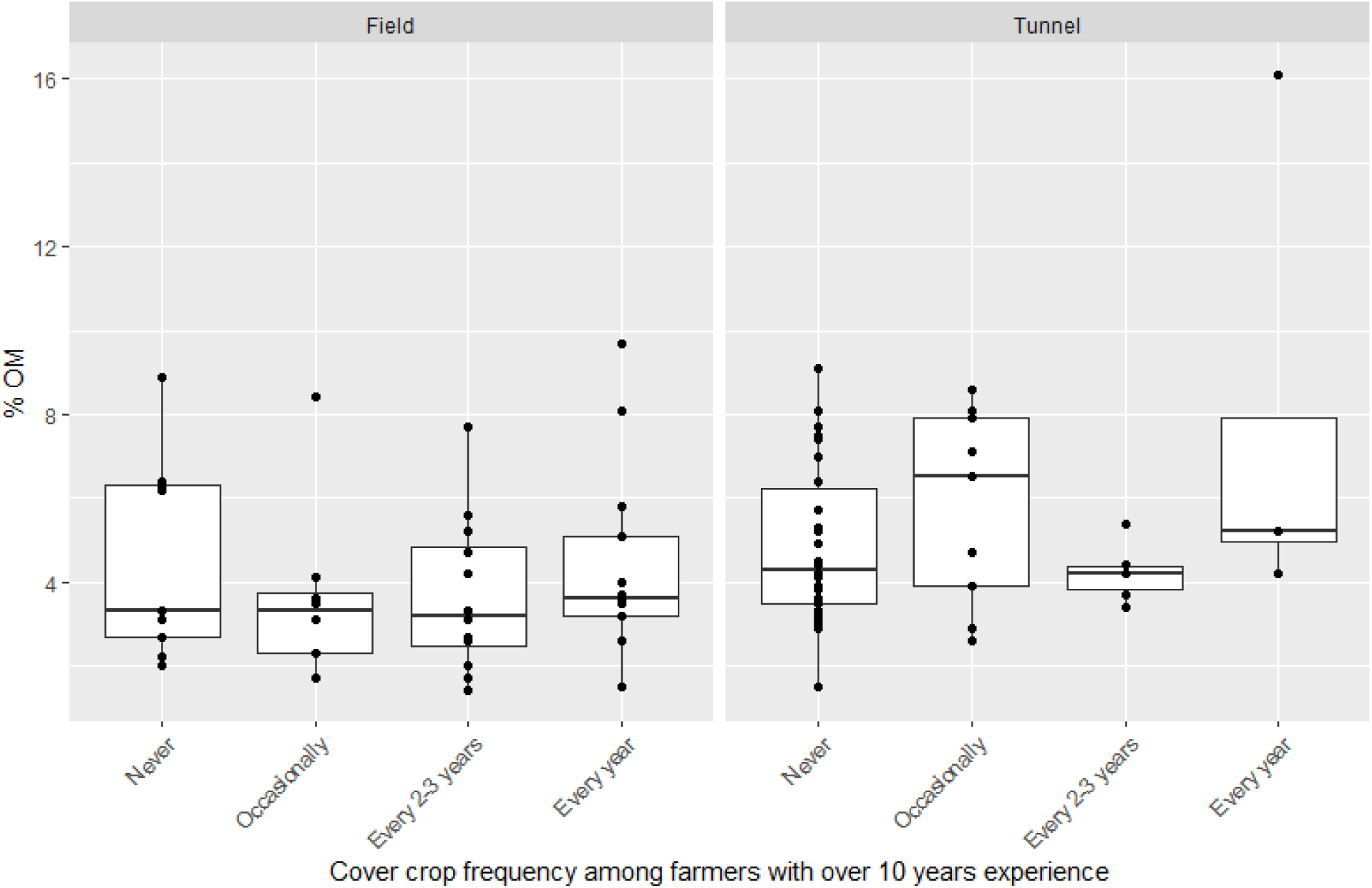
Cover crop frequency and its impact on soil organic matter in fields and high tunnels, filtered to show only farmers with over 10 years of experience.

Finally, more intensive tillage was correlated with lower OM and POX-C, less aggregate stability, and lower soil nitrate and electrical conductivity (Table 6).

Frequency of soil testing was not correlated with any measure of soil health including OM, CEC, POX-C, electrical conductivity, phosphorus, nitrate, bulk density, aggregate stability, infiltration speed, and AMF spore density (Table 5).

#### 3.5.3 Inputs

The use of fresh and composted manure was not correlated with any soil health variable. More frequent use of vegetative compost was correlated with higher OM and aggregate stability, whereas the use of synthetic inputs (both synthetic nitrogen sources like urea and ammonia and synthetic balanced fertilizers like 10-10-10 type products) was correlated with lower OM, CEC, POX-C, more compaction, lower aggregate stability, and higher AMF spore density (Table 7).

**Table 7.** Correlation coefficients and p-values from Spearman’s Rho test for associations between frequency of use of different inputs and soil health metrics.

|  | Fresh manure | Composted manure | Vegetative compost | Synthetic N | Synthetic balanced fertilizers |
| --- | --- | --- | --- | --- | --- |
| **Organic matter** | -0.08<br>p = 0.2890 | 0.03<br>p = 0.6604 | 0.16<br>p = 0.0400 | -0.31<br>p = 0.0001 | -0.36<br>p = 0.0000 |
| **Cation exchange capacity** | -0.10<br>p = 0.1885 | 0.06<br>p = 0.4777 | 0.06<br>p = 0.4240 | -0.28<br>p = 0.0004 | -0.35<br>p = 0.0000 |
| **Permanganate oxidizable carbon (mg C oxidized per kg dry soil)** | -0.04<br>p = 0.5791 | -0.04<br>p = 0.6514 | 0.11<br>p = 0.1592 | -0.19<br>p = 0.0215 | -0.34<br>p = 0.0000 |
| **EC (1:1)** | -0.04<br>p = 0.6097 | 0.07<br>p = 0.3960 | 0.01<br>p = 0.9167 | 0.05<br>p = 0.5321 | -0.08<br>p = 0.3461 |
| **Phosphorus (Bray-P1)** | 0.06<br>p = 0.4264 | 0.02<br>p = 0.8043 | -0.02<br>p = 0.7565 | 0.11<br>p = 0.1695 | 0.15<br>p = 0.0534 |
| **Nitrate** | -0.08<br>p = 0.3258 | 0.06<br>p = 0.4062 | 0.08<br>p = 0.3135 | 0.00<br>p = 0.9693 | -0.01<br>p = 0.8913 |
| **Bulk density** | -0.01<br>p = 0.9056 | -0.04<br>p = 0.6325 | -0.11<br>p = 0.1445 | 0.13<br>p = 0.1017 | 0.18<br>p = 0.0260 |
| **% large soil aggregates** | -0.04<br>p = 0.6454 | 0.03<br>p = 0.6715 | 0.22<br>p = 0.0060 | -0.21<br>p = 0.0109 | -0.35<br>p = 0.0000 |
| **Infiltration rate (mm/hr)** | -0.02<br>p = 0.7792 | 0.13<br>p = 0.1077 | 0.04<br>p = 0.6168 | -0.08<br>p = 0.3768 | -0.17<br>p = 0.0513 |
| **AMF spore density** | 0.12<br>p = 0.1213 | -0.06<br>p = 0.4340 | -0.14<br>p = 0.0858 | 0.30<br>p = 0.0002 | 0.20<br>p = 0.0135 |

## 4 Discussion

### 4.1 High tunnel productivity challenges

Overall, high tunnels in our study had higher organic matter and nutrients than fields, and nutrient concentrations were frequently in excess. This contrasts with the often stated assumption that high tunnel soils may become depleted over time due to longer planting windows and higher yields (DiGiacomo et al., 2023; Nair et al., 2014; Perkus et al., 2022). Our results suggest that while nutrients are certainly taken up by intensive crop production, farmers are replacing them at a faster rate than they deplete them.

Just under half of the high tunnels in our study showed soluble salt accumulation, leading to a classification of “very slightly” or “moderately” saline. This is consistent with farmer-reported concerns and other reports about salt accumulation in high tunnels and associated production challenges (Pierre et al., 2024). However, contrary to the narrative that high tunnels become more saline over time, and that reliance on irrigation water vs. rain water contributes to salt accumulation, we did not see significant relationships between electrical conductivity and farmer experience, years in production, or the pH or alkalinity of irrigation water. Pierre et al. (2024) associated accumulation of mineral salts in tunnels with the combination of high fertilizer application rates and a lack of leaching from rainfall. Electrical conductivity concentrations were positively correlated with soil nitrate concentrations in our study, suggesting high fertilization rates. While farmer reported input application frequency was not correlated with soluble salts, it is likely that the surveyed application frequency variables missed important information about rates of application. For example, a farmer who applies one five-gallon bucket of compost to their high tunnel once per year would have selected the same response in our survey (one application per year) as a farmer who applies two cubic meters of compost once per year. As such, the actual soil nutrient concentrations are likely a better proxy for application rates in our study.

The relationship between soil nitrate and soluble salts also calls into question the practice of flooding high tunnels to manage salt accumulation, a practice commonly recommended by Extension (Brust, 2021; Carpenter et al., 2021; Sánchez, 2023). In Spain where soil-based greenhouse production is widespread, there has been documented nitrate contamination in aquifers. Nitrate leaching from greenhouses in Spain is likely due to overfertilization, particularly with manure, and over-watering, specifically flooding tunnels to leach salts (Thompson et al., 2007). In our study, we estimated that two-thirds of high tunnels and fields have high to moderate potential for nitrate leaching based on soil type, depth to water table, and geology. Rather than focusing on leaching mineral salts from tunnels by flooding them, which in turn may leach nutrients into groundwater, our results suggest that limiting over-fertilization, doing more targeted nutrient applications, and specifically avoiding inputs that are high in salts, may be underutilized strategies for avoiding salt contamination.

Soil pH was above the ideal range of 5.8-7.0 in most high tunnels, and it did not significantly increase with age of high tunnels. This higher pH in high tunnels is important, as plants cannot efficiently utilize nutrients in the soil when the pH is above optimal levels, even if nutrients are abundant in the soil. High soil pH contributed to decreased nitrogen and manganese uptake in high tunnel tomatoes in New York (Reid, 2021). Soil pH should therefore be more closely examined as a source of plant stress in high tunnels. In our study, frequent use of composts and manures did not correlate with pH, though we did not measure actual input amounts. This is consistent with (Sánchez & Ford, 2023b), who did not find a clear correlation between compost use in high tunnels and high pH. Synthetic fertilizers, particularly synthetic nitrogen, have a documented tendency to acidify soil (Barak et al., 1997), and so our finding that more frequent use of synthetic fertilizers correlates with lower soil pH is not surprising. While electrical conductivity and pH were not significantly correlated in our study, the significant correlation between irrigation water with a high alkalinity and pH suggests that water with more dissolved salts is less effective at flushing salts from high tunnels. This should be considered when recommending irrigation-based flooding to flush salts from tunnels.

The final dynamic that may help to explain the problems farmers see in their tunnels is that the soil in many of the high tunnels we visited was extremely dry and had been left bare without a moisture source over the winter. Manure and compost application to the soil can cause water repellency and significantly affect water movement in the soil (Bayad et al., 2020; Pagliari et al., 2011). When high tunnel soil becomes too dry, it can be challenging to re-wet, and irrigation can become a challenge. This dynamic may help to explain the high concentration of fertilizers and salts observed near the soil surface in high tunnels. Educators and researchers in New England have observed that the dry conditions in high tunnels can cause problems for fertilizer solubility and uptake; even soluble fertilizers can remain undissolved in high tunnel soils in dry areas (Hoskins, 2018). Without the natural wetting that comes from rain and snowfall over the winter in Minnesota, high tunnel growers may need to find alternative strategies for maintaining soil moisture over the winter, including irrigated winter cover crops, winter cash crops, or removing plastic.

### 4.2 Nutrient accumulation on diversified vegetable farms

One objective of this research was to confirm the widespread over-abundance of nutrients, particularly phosphorus, on diversified vegetable farms. In both open fields and high tunnels, nutrient concentrations were very high across most farms, and soil phosphorus was particularly high. As with soluble salts, assessing input application frequency was likely not the most robust method of tracking actual inputs since we did not specifically ask growers about rates. Most farmers reported using at least some manure or composted manure every year, generally opting to use a set amount each year, or as much as they can source, rather than calculating the rate based on soil testing (Hoidal, Bugeja, et al., 2025). This, along with the imbalance of nitrogen and phosphorus in organic inputs like compost and manure, is likely to contribute to phosphorus accumulation at many farms.

Using the distance to surface water and soil phosphorus concentrations outlined in the Pennsylvania phosphorus index screening tool (Sharpley et al., 2003), we estimated that only 6 fields and 15 high tunnels in our study had a high risk of having phosphorus runoff reach surface water. However, 35 fields and 36 high tunnels were considered a high-risk or very high-risk distance to surface water, indicating increased potential for runoff. This screening tool has a relatively high phosphorus risk threshold, with a value of 200 ppm (Mehlich-3P), or roughly equal to 74 ppm (Bray-P1) triggering a full analysis. However, at four urban vegetable farms in the Twin Cities, Minnesota, phosphorus became detectable in leachate when soil test phosphorus levels reached about 40 ppm (Bray-1) (Nicklay et al., 2019), which is much lower than the threshold outlined in Sharpley et al., 2023, and much lower than the averages reported in our study. In other analyses of Midwest soils, soil test phosphorus has been shown to be strongly correlated with the amount of phosphorus leaving agricultural fields in surface runoff (Mallarino et al., 2021; Wang et al., 2010).

Globally, agriculture is estimated to contribute approximately 38% of the phosphorus load to freshwater and grey water systems, with the fruit and vegetable sectors each contributing approximately 15% of the agricultural phosphorus budget. This has significant implications for water quality and the ecology of lakes, rivers, and coastal waters (Mekonnen & Hoekstra, 2018). Our results corroborate the global estimates of the contribution of vegetable farmers.

Phosphorus surface runoff is highly variable and depends on factors including soil type, soil moisture, depth of input application, and precipitation. Yet, only very small concentrations of P are needed for a body of water to become eutrophic (Muscutt et al., 1993). Therefore, despite their small footprints on the landscape and further research needed to quantify risk, more education about nutrient management in these systems would be prudent to prevent negative water quality outcomes.

The imbalance of N, P, and K in organic inputs relative to plant needs is well documented, and it is often assumed that growers using organic inputs will over-apply phosphorus to meet nitrogen needs (Möller, 2018). However, our results suggest that some diversified vegetable farms are also applying more nitrogen than plants can efficiently use. Soil tests were conducted in spring before most farmers had applied inputs for the season, and residual nitrate levels were sufficient to support anywhere from 18-59% of crop nitrogen needs, estimated for tomato crops. This was without accounting for organic nitrogen that could become mineralized throughout the growing season, or additional nitrate below 15 cm. This is consistent with findings from high tunnels in Spain, where farmers often apply large quantities of manure and where there is significant leaching of nitrates into groundwater (Thompson et al., 2007). Our risk assessment revealed nitrate leaching risk at two-thirds of the participating farms. In addition to water quality concerns, over-applying nitrogen can cause high nitrate levels in leafy vegetables, which may negatively impact human health and attract more insects (Magdoff & Van Es, 2009). Thus, in addition to the need for outreach and education related to balancing inputs and minimizing nitrate leaching, there may be an additional need to educate farmers about simply over-fertilizing in general. This is especially true given that soil nutrient levels were not correlated with frequency of soil testing.

Finally, both high tunnels and fields also had high calcium concentrations. While many Minnesota soils naturally have high calcium levels due to the presence of calcium carbonate in the bedrock (Johnson et al., 2022), the levels observed in this trial were well above baseline levels for the state, and may pose problems for plant health. Research on high tunnel tomatoes in New York has shown high soil calcium levels may inhibit potassium uptake in tomatoes (Reid, 2021).

### 4.3 Soil health metrics

#### 4.3.1 Organic matter and related metrics

Relative to other farms in the study, the addition of vegetative composts as well as organic certification led to increased OM, with composted manure additions shown not to be predictive of OM increases. The use of synthetic fertilizers correlated with decreases in total soil OM. Maintaining OM in an intensive vegetable production system is a documented challenge (White et al., 2020), and our study highlighted a declining trend in OM, CEC, and POX-C the longer a field had been in vegetable production (Table 6). It’s puzzling that in our study animal manure did not increase OM as others have found increases in soil carbon (Verlinden et al., 2017), especially at high application rates (Evanylo et al., 2008). However, changes in OM with manure are not always detected (Ren et al., 2014) and require a large number of quantified samples (Smith, 2004). It may be that our survey, covering a wide array of farms and soil types, lacked power to detect changes in OM, especially given the lack of measured manure inputs. This finding that vegetative compost is the strongest driver of OM accumulation is consistent with the findings of White et al. (2020), though others have not detected increases in soil carbon with plant derived organic matter inputs such as green manure (Verlinden et al., 2017).

Permanganate oxidizable carbon (POX-C) is an increasingly popular test utilized for soil health assessment, such as in the Cornell Comprehensive Assessment of Soil Health (Moebius-Clune et al., 2016). It is commonly considered to represent an active pool of labile C forms that are likely to be available for microbial metabolism (Culman et al., 2012). Previous research has shown that cover crops may provide more readily decomposable forms of carbon than compost applications (White et al., 2020), and so we had anticipated that active carbon pools might be more closely correlated with cover cropping than total organic matter, but we did not see this in our dataset. This is consistent with research suggesting that POX-C values do not clearly provide improved prediction of soil function compared to SOM values (Koorneef et al., 2024; Liptzin et al., 2022). Additionally, recent work has highlighted the need for caution in interpreting POX-C values, noting that the assay is sensitive not only to labile C forms, but also to lignin and other compounds considered recalcitrant to decomposition (Woodings & Margenot, 2023). Our dataset did not support the assumption that POX-C is more rapidly responsive to management than OM: tillage intensity and frequency of vegetative compost application were notably more predictive of SOM than of POX-C, while none of the measured farm or practice variables were notably more predictive of POX-C than of OM (Tables 6-7). These results suggest that the actionable value of POX-C testing may be limited, and promotion of this test to farmers may not be advisable, especially if the cost is significantly higher than simply assessing organic matter.

We were surprised to see that more frequent cover crop use was correlated with lower OM and slower infiltration, especially given increases in organic matter with vegetative compost additions. Since cover crops may contribute a small and variable amount of biomass carbon to the soil pool relative to cash crops that may be mulched and fertilized, soil changes may proceed slowly and in variable directions as well. Since our study included many beginning farmers, some of the participants had not been farming long enough to see meaningful cover crop benefits. Indeed, when we filtered the data to show only farmers with over 10 years of experience, there was no longer a correlation between cover crop use and lower OM. Our correlation was also developed based on categorical reports of cover crop usage, so we cannot determine the effect of biomass on soil properties in this case, but previous research has found that benefits were biomass-dependent (Finney et al., 2016). A potential explanation for slower infiltration with higher cover crop frequency could be more passes by foot or equipment, which could compact soil. Farmers with more compact soil may also be more likely to try cover crops as a mitigation strategy than farmers who do not struggle to manage compaction. On the positive side for cover crops, the negative correlations of soil Bray-P and nitrate with cover crop use frequency suggests that cover crops may be effectively pulling in plant-available nutrients and potentially keeping them in organic forms. This is critical for cover crops to achieve proposed water quality benefits (Lewandowski & Cates, 2023), and a first step in holding nutrients for later cash crop nutrient supply in an organic system. Farmers may also be less likely to over-apply fertilizers if they are using cover crops as part of their fertility strategy.

#### 4.3.2 Physical measures of soil health

Overall physical soil health parameters, including infiltration, were lower among more experienced farmers and in fields with a longer history of vegetable production, highlighting the challenge of maintaining soil health in intensive vegetable production systems. Compaction from foot traffic and small machinery is under-studied relative to row crop compaction, but literature from turf systems suggests that even lightweight traffic can increase surface hardness and bulk density (Braun et al., 2022). Physical properties are linked to water quality outcomes; specifically, higher aggregate stability reduces risk and rates of surface runoff and nutrient loss (Barthès & Roose, 2002). Since most vegetable fields and tunnels were over-fertilized, these systems are at particularly high risk for surface nutrient losses. As with OM, the only input positively correlated with aggregate stability was vegetative compost, but compost had no impact on bulk density or infiltration.

Soil aggregation can vary widely across soil type and climate (Ciric et al., 2012; Liu et al., 2021), yet we were able to see a significant correlation between reducing tillage intensity and improved aggregate stability. This is consistent with research identifying tillage as a major driver of aggregate stability in cropping systems (Six et al., 2000).

Although aggregation is strongly theoretically connected to infiltration rates (Crawford, 1994), in practice it is common for infiltration to lag in sensitivity behind soil aggregation (Basset et al., 2023; Blair et al., 2024; Garg et al., 2025). Infiltration is so variable in the field that detecting differences between treatments is challenging regardless of methods (Schott et al., 2023).

Aggregation is somewhat more responsive to various treatments across soil types than infiltration or bulk density (Basset et al., 2023), and our study confirms that this parameter is sensitive to management across a wide variety of crops, soils, and management strategies (Douds et al., 1997; Zhang et al., 2021).

#### 4.3.4 Arbuscular mycorrhizal fungi

AMF spore counts did not significantly correlate with most soil health practices, but they were positively correlated with more frequent use of synthetic inputs, and with conventionally managed (non-organic) farms. This result aligns with previous observations of high soil fertility rates correlated with high spore numbers. Previous authors have hypothesized that in the presence of readily available nutrients, plants are less likely to signal for AMF spore germination (Douds et al., 1997; Zhang et al., 2021). To confirm this hypothesis, it would be useful to measure AMF colonization in plants growing in the respective soil indicative of active symbiosis.

## 5 Conclusions

Our study of 100 high tunnels and 100 fields at small-scale vegetable farms in Minnesota confirmed that diversified vegetable farmers face a unique set of soil health challenges. High tunnels in our study were generally high in organic matter and nutrients but tended to have a high soil pH, and nearly half of the high tunnels had problems with soluble salts. More nitrates and more frequent use of synthetic fertilizers were associated with higher soluble salt concentrations, and alkaline irrigation water seemed to be a significant driver of soil pH increases. This finding highlights the importance of careful nutrient management in high tunnels, and the need for farmers to focus on sourcing inputs with low salt concentrations.

In both high tunnel and field environments, nutrient concentrations were very high, and phosphorus tended to be far in excess of the recommended range. Soil nitrate concentrations were high in both environments considering the time of year (pre-field preparation for most participants). Over-fertilization on these farms likely poses a threat to both surface and groundwater, and this study highlights the need for more targeted nutrient management strategies on diversified vegetable farms.

Vegetable production practices had a mixed impact on most of the soil health indicators in our study. There was a general trend towards decreasing soil health metrics the longer a field had been in vegetable production, and among farmers with more experience. However, strategies like reducing tillage, applying vegetative compost, and using organic production methods increased organic matter and aggregation. These results suggest that while vegetable production may be hard on soils, conservation practices can significantly improve the overall health of the soil in these intensive production systems.

## Abbreviations

CEC: cation exchange capacity
OM: organic matter
NRCS: Natural Resource Conservation Service
POX-C: permanganate oxidizable carbon

## Notes

### Competing Interest Statement

The authors have declared no competing interest.

### Funding

- Minnesota Agricultural Experiment Station: Rapid Response Fund

## REFERENCES

Bailey, P., & Kagan, A. (2020). *Emerging farmers in Minnesota* (Legislative Report Minnesota Publication 20-0237). Minnesota Department of Agriculture.

Barak, P., Jobe, B. O., Krueger, A. R., Peterson, L. A., & Laird, D. A. (1997). Effects of long-term soil acidification due to nitrogen fertilizer inputs in Wisconsino title found]. Plant and Soil, 197(1), 61–69. 10.1023/A:1004297607070

Barthès, B., & Roose, E. (2002). Aggregate stability as an indicator of soil susceptibility to runoff and erosion; validation at several levels. CATENA, 47(2), 133–149. 10.1016/S0341-8162(01)00180-1

Basset, C., Abou Najm, M., Ghezzehei, T., Hao, X., & Daccache, A. (2023). How does soil structure affect water infiltration? A meta-data systematic review. Soil and Tillage Research, 226, 105577. 10.1016/j.still.2022.105577

Bayad, M., Chau, H. W., Trolove, S., Moir, J., Condron, L., & Bouray, M. (2020). The Relationship between Soil Moisture and Soil Water Repellency Persistence in Hydrophobic Soils. Water, 12(9), 2322. 10.3390/w12092322

Blair, H. K., Gutknecht, J. L., Jelinski, N. A., Lewandowski, A. M., Fisher, B. A., & Cates, A. M. (2024). Nature versus nurture: Quantifying the effects of management, region, and hillslope position on soil health indicators in an on-farm survey in Minnesota. Soil Science Society of America Journal, 88(6), 2135–2155. 10.1002/saj2.20739

Braun, R. C., Bremer, D. J., & Hoyle, J. A. (2022). Simulated traffic on turfgrasses during drought stress: II. Soil moisture, soil compaction, and rooting. International Turfgrass Society Research Journal, 14(1), 516–527. 10.1002/its2.62

Bruce, A. B., Maynard, E. T., & Farmer, J. R. (2019). Farmers’ Perspectives on Challenges and Opportunities Associated with Using High Tunnels for Specialty Crops. HortTechnology, 29(3), 290–299. 10.21273/HORTTECH04258-18

Brust, G. (2021, September). High Soluble Salts a Problem in Some High Tunnels. In University of Maryland Extension. https://extension.umd.edu/resource/high-soluble-salts-problem-some-high-tunnels/

Camberato, J., & Nielsen, B. (2017, June). Soil Sampling to Assess Current Soil N Availability. Purdue Agronomy Corny News. https://www.agry.purdue.edu/ext/corn/news/timeless/assessavailablen.html

Carpenter, J., Basden, T., Rayburn, E., Jett, L., & McDonald, L. (2021, February). Salinity Management in High Tunnel Systems. In West Virginia Univeristy Extension. https://extension.wvu.edu/natural-resources/soil-water/salinity-management-in-high-tunnel-systems

Ciric, V., Manojlovic, M., Nesic, L., & Belic, M. (2012). Soil dry aggregate size distribution: Effects of soil type and land use. Journal of Soil Science and Plant Nutrition, ahead, 0–0. 10.4067/S0718-95162012005000025

Cox, D. (1995, August). Water Quality: pH and Alkalinity. UMass Extension Greenhouse Crops and Floriculture Program. https://ag.umass.edu/greenhouse-floriculture/fact-sheets/water-quality-ph-alkalinity

Crawford, J. W. (1994). The relationship between structure and the hydraulic conductivity of soil. European Journal of Soil Science, 45(4), 493–502. 10.1111/j.1365-2389.1994.tb00535.x

Culman, S., Mann, M., Sharma, S., Saeed, M. T., Fulford, A., Lindsey, L., Brooker, A., Dayton, L., Eugene, B., Warden, R., Steinke, K., Camerato, J., & Joern, B. (2019, June 14). Converting between Mehlich-3, Bray P, and Ammonium Acetate Soil Test Values. Ohio State University Extension. https://ohioline.osu.edu/factsheet/anr-75

Culman, S. W., Snapp, S. S., Freeman, M. A., Schipanski, M. E., Beniston, J., Lal, R., Drinkwater, L. E., Franzluebbers, A. J., Glover, J. D., Grandy, A. S., Lee, J., Six, J., Maul, J. E., Mirksy, S. B., Spargo, J. T., & Wander, M. M. (2012). Permanganate Oxidizable Carbon Reflects a Processed Soil Fraction that is Sensitive to Management. Soil Science Society of America Journal, 76(2), 494–504. 10.2136/sssaj2011.0286

DeLong, A., Swisher, M. E., Chase, C. A., Irani, T., & Ruiz-Menjivar, J. (2023). The Roots of First-Generation Farmers: The Role of Inspiration in Starting an Organic Farm. Land, 12(6), 1169. 10.3390/land12061169

DiGiacomo, G., Gieske, M., Grossman, J., Jacobsen, K., Peterson, H., & Rivard, C. (2023). Economic trade-offs: Analysis of hairy vetch (*Vicia villosa*) cover crop use in organic tomato (*Solanum lycopersicum* L.) high tunnel systems across multiple regions. Renewable Agriculture and Food Systems, 38, e10. 10.1017/S1742170523000029

Douds, D. D., Galvez, L., Franke-Snyder, M., Reider, C., & Drinkwater, L. E. (1997). Effect of compost addition and crop rotation point upon VAM fungi. Agriculture, Ecosystems & Environment, 65(3), 257–266. 10.1016/S0167-8809(97)00075-3

Evanylo, G., Sherony, C., Spargo, J., Starner, D., Brosius, M., & Haering, K. (2008). Soil and water environmental effects of fertilizer-, manure-, and compost-based fertility practices in an organic vegetable cropping system. Agriculture, Ecosystems & Environment, 127(1–2), 50–58. 10.1016/j.agee.2008.02.014

Finney, D. M., White, C. M., & Kaye, J. P. (2016). Biomass Production and Carbon/Nitrogen Ratio Influence Ecosystem Services from Cover Crop Mixtures. Agronomy Journal, 108(1), 39–52. 10.2134/agronj15.0182

Garg, A., Kwakye, S., Cates, A., Peterson, H., LaBine, K., Olson, G., & Sharma, V. (2025). Integrated soil health management influences soil properties: Insights from a US Midwest study. Geoderma, 455, 117214. 10.1016/j.geoderma.2025.117214

Gerdemann, J. W., & Nicolson, T. H. (1963). Spores of mycorrhizal Endogone species extracted from soil by wet sieving and decanting. Transactions of the British Mycological Society, 46(2), 235–244. 10.1016/S0007-1536(63)80079-0

Grigal, D. F. (1973). *Note on the Hydrometer Method of Particle-Size Analysis* (245; Minnesota Forestry Research Notes). University of Minnesota Digital Conservancy. https://hdl.handle.net/11299/58296

Hoidal, N., Bugeja, S. M., Lindenfelser, E., & Pagliari, P. H. (2025). Soil Health Practices and Decision Drivers on Diversified Vegetable Farms in Minnesota. Sustainability, 17(3), 1192. 10.3390/su17031192

Hoidal, N., Fernandez, A., Kubovcik, K., Berry, K., Andreasen, A., Ispache, D., & Grossman, J. (2024). Emerging specialty crop farmer perspectives and educational needs related to soil health and nutrient management in the Upper Midwest. Renewable Agriculture and Food Systems, 39, e25. 10.1017/S1742170524000139

Hoidal, N., Rosen, C., Pagliari, P., & Fernandez, A. (2025). Soil health and nutrient management in high tunnels. University of Minnesota Extension. https://extension.umn.edu/nutrient-management-specialty-crops/soil-health-and-nutrient-management-high-tunnels

Hoskins, B. (2018, February 15). High tunnel soil management update. UMass Extension Vegetable Notes; 30(2). https://ag.umass.edu/sites/ag.umass.edu/files/newsletters/february_15_2018_vegetable_notes.pdf

Johnson, L., Bartsch, W., Hudak, G., Davenport, M., Johnson, K., Nixon, K., Reed, J., & team, A. (2022). Minnesota Natural Resource Atlas: Online mapping tools and data for natural resource planning, management, and research in Minnesota. In Natural Resources Research Institute, University of Minnesota Duluth.

Klodd, A., & Hoidal, N. (2019). Needs Assessment of Minnesota Fruit and Vegetable Producers. University of Minnesota Extension. https://conservancy.umn.edu/handle/11299/208753

Konert, M., & Vandenberghe, J. (1997). Comparison of laser grain size analysis with pipette and sieve analysis: A solution for the underestimation of the clay fraction. Sedimentology, 44(3), 523–535. 10.1046/j.1365-3091.1997.d01-38.x

Koorneef, G. J., Pulleman, M. M., Comans, R. Nj., Van Rijssel, S. Q., Barré, P., Baudin, F., & De Goede, R. Gm. (2024). Assessing soil functioning: What is the added value of soil organic carbon quality measurements alongside total organic carbon content? Soil Biology and Biochemistry, 196, 109507. 10.1016/j.soilbio.2024.109507

Lamont, W. J. (2009). Overview of the Use of High Tunnels Worldwide. HortTechnology, 19(1), 25–29. 10.21273/HORTTECH.19.1.25

Lewandowski, A. M., & Cates, A. (2023). Connecting soil health and water quality in agricultural landscapes. Journal of Environmental Quality, 52(3), 412–421. 10.1002/jeq2.20390

Liptzin, D., Norris, C. E., Cappellazzi, S. B., Bean, G. M., Cope, M., Greub, K. L. H., Rieke, E. L., Tracy, P. W., Aberle, E., Ashworth, A., Bañuelos Tavarez, O., Bary, A. I., Baumhardt, R. L., Borbón Gracia, A., Brainard, D. C., Brennan, J. R., Briones Reyes, D., Bruhjell, D., Carlyle, C. N., … Honeycutt, C. W. (2022). An evaluation of carbon indicators of soil health in long-term agricultural experiments. Soil Biology and Biochemistry, 172, 108708. 10.1016/j.soilbio.2022.108708

Liu, X., Wu, X., Liang, G., Zheng, F., Zhang, M., & Li, S. (2021). A global meta-analysis of the impacts of no-tillage on soil aggregation and aggregate-associated organic carbon. Land Degradation & Development, 32(18), 5292–5305. 10.1002/ldr.4109

Magdoff, F., & Van Es, H. (2009). Building soils for better crops: Sustainable soil management (3rd ed). Sustainable Agriculture Research and Education (SARE).

Mallarino, A. P., Haq, M. U., & Jones, J. D. (2021). Report to the Iowa nutrient research center, project 2017-05. Amounts and forms of dissolved phosphorus lost with surface runoff as affected by phosphorus management and soil conservation practices.

Mekonnen, M. M., & Hoekstra, A. Y. (2018). Global Anthropogenic Phosphorus Loads to Freshwater and Associated Grey Water Footprints and Water Pollution Levels: A High-Resolution Global Study. Water Resources Research, 54(1), 345–358. 10.1002/2017WR020448

Miller, B. A., & Schaetzl, R. J. (2012). Precision of Soil Particle Size Analysis using Laser Diffractometry. Soil Science Society of America Journal, 76(5), 1719–1727. 10.2136/sssaj2011.0303

Minnesota Department of Agriculture. (2025). Vulnerable Groundwater Area Map [Dataset]. https://www.mda.state.mn.us/chemicals/fertilizers/nutrient-mgmt/nitrogenplan/mitigation/wrpr/wrprpart1/vulnerableareamap

Minnesota Department of Natural Resources. (2016). Water-Table Elevation and Depth to Water Table, Minnesota Hydrogeology Atlas series HG-03. [Dataset]. Minnesota Geospatial Commons. https://gisdata.mn.gov/dataset/geos-hydrogeology-atlas-hg03

Minnesota Department of Natural Resources. (2019). National Wetland Inventory for Minnesota [Dataset]. Minnesota Geospatial Commons. https://gisdata.mn.gov/dataset/water-nat-wetlands-inv-2009-2014

Minnesota Geospatial Information Office. (2012). Lidar Elevation Data for Minnesota: 2008-2012 [Dataset]. Minnesota Geospatial Information Office. https://www.mngeo.state.mn.us/chouse/elevation/lidar_2008-2012.html

Moebius-Clune, B. D., Moebius-Clune, D. J., Gugino, B. K., Idowu, O. J., Schindelbeck, R. R., Ristow, A. J., van Es, H. M., Thies, J. E., Shayler, H. A., McBride, M. B., Kurtz, K. S. M., Wolfe, D. W., & Abawi, G. S. (2016). Comprehensive Assessment of Soil Health – The Cornell Framework, Edition 3. Cornell University.

Möller, K. (2018). Soil fertility status and nutrient input–output flows of specialised organic cropping systems: A review. Nutrient Cycling in Agroecosystems, 112(2), 147–164. 10.1007/s10705-018-9946-2

Muscutt, A. D., Harris, G. L., Bailey, S. W., & Davies, D. B. (1993). Buffer zones to improve water quality: A review of their potential use in UK agriculture. Agriculture, Ecosystems and Environment, 45, 59–77.

Nair, A., Carpenter, B. H., Tillman, J. L., & Jokela, D. L. (2014). Integrating cover crops in high tunnel crop production. 2009. Iowa State Research Farm Progress Reports.

National Oceanic and Atmospheric Administration. (2022, October). Hydrologic soils group. National Oceanic and Atmospheric Administration Office for Coastal Management. https://coast.noaa.gov/data/digitalcoast/pdf/qnspect-hydrologic-soils.pdf

National Sustainable Agriculture Coalition. (2016). Environmental Quality Incentives Program. https://sustainableagriculture.net/publications/grassrootsguide/conservation-environment/environmental-quality-incentives-program/

Natural Resources Conservation Service. (2001). Soil quality test kit guide. United States Department of Agriculture Agricultural Research Service, Soil Quality Institute. https://www.nrcs.usda.gov/sites/default/files/2022-10/Soil%20Quality%20Test%20Kit%20Guide.pdf

Nicklay, J., Cadieux, V., Jelinski, N., LaBine, K., Rogers, M., & Small, C. (2019). Initial trends in ecosystem service metrics of urban agriculture in Minneapolis/St. Paul, MN. Oral Session Presented at the ASA-CSSA-SSSA International Annual Meeting.

Pagliari, P. H., Flores-Mangual, M. L., Lowery, B., Weisenberger, D. G., & Laboski, C. A. M. (2011). Manure-Induced Soil-Water Repellency. Soil Science, 176(11), 576–581. 10.1097/SS.0b013e3182316c7e

Perkus, E. A., Grossman, J. M., Pfeiffer, A., Rogers, M. A., & Rosen, C. J. (2022). Exploring Overwintered Cover Crops as a Soil Management Tool in Upper-midwest High Tunnels. HortScience, 57(2), 171–180. 10.21273/HORTSCI15987-21

Pierre, J. F., Jacobsen, K. L., Wszelaki, A., Butler, D., Velandia, M., Woods, T., Sideman, R., Grossman, J., Coolong, T., Hoskins, B., Da Silva, A. L. B. R., Ginakes, P., Kleinhenz, M., Zhao, X., Rivard, C., & Rudolph, R. E. (2024). Sustaining Soil Health in High Tunnels: A Paradigm Shift toward Soil-centered Management. HortTechnology, 34(5), 594–603. 10.21273/HORTTECH05460-24

R Core Team. (2022). R: A language and environment for statistical computing. (Version v4.2.2) [Computer software]. R Foundation for Statistical Computing. https://www.R-project.org/

Reid, J. (2021, February). The unseen elephant in the room: Why and how to manage calcium in high tunnel soils [Webinar]. UMass Amherst Extension Vegetable Program High Tunnel Fertility Research Update. https://www.youtube.com/watch?v=TzXbqoticu0

Ren, T., Wang, J., Chen, Q., Zhang, F., & Lu, S. (2014). The Effects of Manure and Nitrogen Fertilizer Applications on Soil Organic Carbon and Nitrogen in a High-Input Cropping System. PLoS ONE, 9(5), e97732. 10.1371/journal.pone.0097732

Rosen, C., & Eliason, R. (2005). Nutrient Management for Commercial Fruit & Vegetable Crops in Minnesota. Unviersity of Minnesota Extension.

Ruch, B., Hefner, M., & Sradnick, A. (2023). Excessive Nitrate Limits the Sustainability of Deep Compost Mulch in Organic Market Gardening. Agriculture, 13(5), 1080. 10.3390/agriculture13051080

Rudisill, M. A., Bordelon, B. P., Turco, R. F., & Hoagland, L. A. (2015). Sustaining Soil Quality in Intensively Managed High Tunnel Vegetable Production Systems: A Role for Green Manures and Chicken Litter. HortScience, 50(3), 461–468. 10.21273/HORTSCI.50.3.461

Sánchez, E. (2023, March). Dealing with High Soluble Salt Levels in High Tunnels. In Penn State Extension. https://extension.psu.edu/dealing-with-high-soluble-salt-levels-in-high-tunnels

Sánchez, E., & Ford, T. (2023a). High Tunnel Soil Test Report: Soil Nutrient Levels. In Penn State Extension. https://extension.psu.edu/high-tunnel-soil-test-report-soil-nutrient-levels

Sánchez, E., & Ford, T. (2023b). High Tunnel Soil Test Report: Soil pH. In Penn State Extension. https://extension.psu.edu/high-tunnel-soil-test-report-soil-ph

Schott, L. R., Yost, J. L., Kruger, K., Leytem, A. B., & Dungan, R. S. (2023). Assessment of Infiltration Methodologies for Calcareous Silty Soils in Idaho’s Magic Valley. Soil Erosion Research Under a Changing Climate, January 8-13, 2023, Aguadilla, Puerto Rico, USA. Soil Erosion Research Under a Changing Climate, January 8-13, 2023, Aguadilla, Puerto Rico, USA. 10.13031/soil.23039

Sharpley, A. N., Weld, J. L., Beegle, D. B., Kleinman, P. J. A., Gburek, W. J., Moore, P. A., & Mullins, G. (2003). Development of phosphorus indices for nutrient management planning strategies in the United States. Journal of Soil and Water Conservation, 58(3), 137.

Shrestha, P., Small, G. E., & Kay, A. (2020). Quantifying nutrient recovery efficiency and loss from compost-based urban agriculture. PLOS ONE, 15(4), e0230996. 10.1371/journal.pone.0230996

Six, J., Elliott, E. T., & Paustian, K. (2000). Soil macroaggregate turnover and microaggregate formation: A mechanism for C sequestration under no-tillage agriculture. Soil Biology and Biochemistry, 32(14), 2099–2103. 10.1016/S0038-0717(00)00179-6

Small, G., Shrestha, P., Metson, G. S., Polsky, K., Jimenez, I., & Kay, A. (2019). Excess phosphorus from compost applications in urban gardens creates potential pollution hotspots. Environmental Research Communications, 1(9), 091007. 10.1088/2515-7620/ab3b8c

Smith, P. (2004). How long before a change in soil organic carbon can be detected? Global Change Biology, 10(11), 1878–1883. 10.1111/j.1365-2486.2004.00854.x

Soil Survey Staff. (2022). Kellogg Soil Survey Laboratory Methods Manual (oil Survey Investigations Report No. 42 Version 6.0).

Sullivan, C. (2015). The Minnesota farmer: Demographic trends and relevant laws (A Changing Minnesota). House research department and Minnesota state demographic center. https://www.house.mn.gov/hrd/pubs/minnfarmer.pdf

Thompson, R. B., Martínez-Gaitan, C., Gallardo, M., Giménez, C., & Fernández, M. D. (2007). Identification of irrigation and N management practices that contribute to nitrate leaching loss from an intensive vegetable production system by use of a comprehensive survey. Agricultural Water Management, 89(3), 261–274. 10.1016/j.agwat.2007.01.013

United States. (2012). U.S. Census of Agriculture. Library of Congress. https://www.loc.gov/item/lcwaN0022771/

United States. (2022). U.S. Census of Agriculture. Library of Congress. https://www.loc.gov/item/lcwaN0022771/

University of Minnesota. (2024a). University of Minnesota Research Analytical Laboratory: Major Equipment. https://ral.cfans.umn.edu/instruments-methods/major-equipment

University of Minnesota. (2024b). University of Minnesota Research Analytical Laboratory: Method References. https://ral.cfans.umn.edu/instruments-methods/method-references

Verlinden, S., McDonald, L., Kotcon, J., & Childs, S. (2017). Long-term Effect of Manure Application in a Certified Organic Production System on Soil Physical and Chemical Parameters and Vegetable Yields. HortTechnology, 27(2), 171–176. 10.21273/HORTTECH03348-16

Wang, Y. T., Zhang, T. Q., Hu, Q. C., Tan, C. S., Halloran, I. P. O., Drury, C. F., Reid, D. K., Ma, B. L., Ball-Coelho, B., Lauzon, J. D., Reynolds, W. D., & Welacky, T. (2010). Estimating Dissolved Reactive Phosphorus Concentration in Surface Runoff Water from Major Ontario Soils. Journal of Environmental Quality, 39(5), 1771–1781. 10.2134/jeq2009.0504

Weil, R. R., Islam, K. R., Stine, M. A., Gruver, J. B., & Samson-Liebig, S. E. (2003). Estimating active carbon for soil quality assessment: A simplified method for laboratory and field use. American Journal of Alternative Agriculture, 18(1), 3–17. 10.1079/AJAA200228

White, K. E., Brennan, E. B., Cavigelli, M. A., & Smith, R. F. (2020). Winter cover crops increase readily decomposable soil carbon, but compost drives total soil carbon during eight years of intensive, organic vegetable production in California. PLOS ONE, 15(2), e0228677. 10.1371/journal.pone.0228677

Wilson, M. (2021). Manure characteristics. In University of Minnesota Extension. https://extension.umn.edu/manure-management/manure-characteristics

Woodings, F. S., & Margenot, A. J. (2023). Revisiting the permanganate oxidizable carbon (POXC) assay assumptions: POXC is lignin sensitive. Agricultural & Environmental Letters, 8(1), e20108. 10.1002/ael2.20108

Yoder, R. E. (1936). A Direct Method of Aggregate Analysis of Soils and a Study of the Physical Nature of Erosion Losses^1^. Agronomy Journal, 28(5), 337–351. 10.2134/agronj1936.00021962002800050001x

Zhang, S., Luo, P., Yang, J., Irfan, M., Dai, J., An, N., Li, N., & Han, X. (2021). Responses of Arbuscular Mycorrhizal Fungi Diversity and Community to 41-Year Rotation Fertilization in Brown Soil Region of Northeast China. Frontiers in Microbiology, 12, 742651. 10.3389/fmicb.2021.742651

